# Data Representation Bias and Conditional Distribution Shift Drive Predictive Performance Disparities in Multi-Population Machine Learning

**DOI:** 10.1101/2025.06.18.658431

**Authors:** Sandeep Kumar, Yan Cui

## Abstract

Machine learning frequently encounters challenges when applied to population-stratified datasets, where data representation bias and data distribution shifts substantially impact model performance and generalizability across different population groups. These challenges are well illustrated in the context of polygenic prediction for diverse ancestry groups, and the underlying mechanisms are broadly applicable to machine learning with population-stratified data across domains. Using synthetic genotype-phenotype datasets representing five continental populations, we evaluate three approaches for utilizing population-stratified data, mixture learning, independent learning, and transfer learning, to systematically investigate how data representation bias and distribution shifts influence multi-population machine learning. Our results show that conditional distribution shifts, in combination with data representation bias, significantly influence machine learning performance across diverse populations and the effectiveness of transfer learning as a disparity mitigation strategy, while the effect of marginal distribution shifts is limited. The joint effects of data representation bias and distribution shifts demonstrate distinct patterns under different multi-population machine learning approaches, providing critical insights for the development of effective and equitable machine learning models for population-stratified data.

## Introduction

Population stratification, the presence of distinct subgroups within a population that differ systematically in characteristics such as age, sex, genetic ancestry, or socioeconomic status, significantly impacts machine learning applications across various domains [1–3]. It is of crucial importance to ensure equitable machine learning across diverse groups by identifying and mitigating model performance disparities [4]. In genomics, for example, polygenic prediction of diseases has great potential to advance genomic medicine by utilizing large-scale genomic data to improve the accuracy of disease risk assessments and enable personalized prevention and treatment strategies [5, 6]; however, the effectiveness of polygenic prediction is often compromised by representation bias of genomic data across different ancestry groups. Most genomic research has been conducted on populations of primarily European descent, leading to less accurate risk predictions for individuals from underrepresented groups [7–18]. Compounding the challenge of representation bias, data distribution shifts across different ancestry groups further complicate and significantly impact model performance and generalizability [19–21]. Addressing this challenge requires diversifying genomic data and developing machine learning strategies to enable that polygenic prediction benefits are more equitably applicable, thereby improving the effectiveness of genomic medicine for all populations [20, 22].

Cross-ancestry generalizability in polygenic models has been enhanced through approaches that recalibrate genetic effect sizes or adjust model sparsity and shrinkage parameters across ancestry groups [18, 23–29]. However, conventional polygenic prediction methods are predominantly based on linear additive modeling frameworks [30], which can limit their capacity to learn and transfer complex representations across ancestry groups. More recently, machine learning approaches capable of modeling complex nonlinear patterns have been applied to genomic disease prediction [31–34], and these models have outperformed conventional polygenic prediction methods in many studies [19, 35–42]. Machine learning models such as neural networks possess substantially greater model capacity for capturing complex, non-linear relationships and are particularly well suited for transfer learning [43]. Recent studies have shown that transfer learning is effective in improving omics-based clinical predictions for data-disadvantaged populations [44–47]. Notably, transfer learning improved predictive accuracy for disadvantaged populations without compromising performance for other population groups, thus achieving a generally desirable Pareto improvement [19, 48].

This study investigates a fundamental machine learning question that extends beyond polygenic prediction to any domain involving population-stratified data: how data representation bias and distribution shifts shape the landscape of multi-population machine learning performance. We use synthetic genotype-phenotype datasets as a controlled experimental platform because genomic data across ancestry groups provide a well-characterized, biologically grounded instantiation of these challenges, enabling precise manipulation of marginal distribution shifts (allele frequency differences), conditional distribution shifts (genotype-phenotype relationship differences), and representation bias independently. Specifically, we address two questions: (1) How do conditional and marginal distribution shifts influence machine learning model performance across different population groups? (2) Under what conditions can transfer learning effectively improve performance for data-disadvantaged populations? By systematically exploring the parameter space that determines data representation bias and distribution shift, we identify regions where model performance disparities emerge and where transfer learning mitigates them. Our results reveal the distinct effects of conditional and marginal distribution shifts on multi-population machine learning across a broad range of conditions, providing generalizable insights for addressing performance disparities in population-stratified machine learning.

## Methods

### Marginal and conditional distribution shifts in multi-population machine learning

A supervised machine learning task 𝒯 typically consists of three essential elements: an input feature matrix 𝑋, an outcome variable 𝑌, and a function 𝑓 mapping the features to the outcome. The task is to learn this function from training data. From a probabilistic perspective, the function 𝑓 can be expressed as 𝑃(𝑌|𝑋). It is generally assumed that each data point is independently drawn from a single joint distribution 𝑃(𝑌, 𝑋) . However, in practice, this assumption rarely holds, especially in the context of machine learning with population-stratified data because the distributions often vary across population groups. Considering 𝑃(𝑌, 𝑋) = 𝑃(𝑌|𝑋)𝑃(𝑋), shifts in either the marginal distribution 𝑃(𝑋) or the conditional distribution 𝑃(𝑌|𝑋) can lead to changes in the joint distribution 𝑃(𝑌, 𝑋), thus impacting machine learning performance. A marginal distribution shift involves changes in the feature space, a conditional distribution shift reflects changes in the relationship between features and outcomes, altering the mapping function 𝑓 that the model aims to learn. While both types of shifts pose challenges for machine learning, conditional distribution shifts more profoundly impact the predictive performance of machine learning models [49, 50].

In the context of polygenic prediction, 𝑋 represents genotype data for a set of genetic variants and 𝑌 is the associated phenotype value for prediction. The marginal distribution 𝑃(𝑋) represents the allele frequencies of genetic variants. The conditional distribution 𝑃(𝑌|𝑋) captures the genotype-phenotype relationship, which is determined by the effect sizes of genetic variants. Genetic architectures of many phenotypes, primarily characterized by allele frequencies and effect sizes of genetic variants, differ among ancestry groups [51–54], resulting in marginal and conditional distribution shifts.

### Multi-population machine learning schemes

Properly utilizing population-stratified data that are subject to representation bias and distribution shifts is critical to reducing machine learning performance disparities across diverse population groups. We have classified different approaches for machine learning with population-stratified data, such as multi-ancestry genotype-phenotype data [20, 44]. *Mixture learning* combines data from all ancestry groups to train a single model. *Independent learning* trains separate models for each group. *Transfer learning* initially trains a model on EUR data (source domain) and subsequently uses DDP data (target domain) for fine-tuning or domain adaptation. These three approaches represent fundamentally different strategies for handling population-stratified data. We evaluate them not as competing methods for maximizing predictive accuracy, but as controlled experimental conditions that allow us to observe how representation bias and distribution shifts manifest differently depending on how multi-population data are utilized.

### Synthetic data with controlled data representation bias and distribution shifts

We utilized a synthetic genotype dataset of 600,000 individuals across five ancestry groups representing the five super-populations in the 1000 Genomes Project [55]: African (AFR), Admixed American (AMR), East Asian (EAS), European (EUR), and South Asian (SAS), each with 120,000 subjects [29, 56]. We retrieved the genotype data from this dataset and generated phenotype data for our machine learning experiments. Each synthetic dataset comprises genotype-phenotype data from two ancestry groups: Europeans (EUR) and a data-disadvantaged non-EUR group (DDP; one of AFR, AMR, EAS, or SAS).

Phenotype values were generated using a statistical model: *Y = G×β + ɛ*

- *G*: Genotype matrix of dimension *N*×*M*, where *N* is the number of individuals (across EUR and DDP) and *M* is the number of SNPs, with genotypes coded as 0, 1, or 2 copies of the effect allele.
- *β*: Vector of effect sizes for the SNPs, sampled from a multivariate normal distribution that induces a specified cross-ancestry genetic correlation ρ. For each SNP 𝑗, the pair of ancestry-specific effects (𝛽_𝑗,EUR_, 𝛽_𝑗,DDP_) was drawn from a bivariate normal 𝒩(0, 𝑅), 𝑅 = 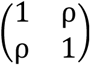, so that the correlation of effects between ancestries is ρ. The variance of genetic component *G×*β was scaled to match the target heritability (*h^2^*).
- *ɛ*: Environmental noise, sampled from a normal distribution 𝒩(0,1− *h^2^*).

When generating binary phenotype values, the continuous phenotype values are dichotomized into case-control categories based on the case to control ratio.

All SNPs in our simulation are treated as directly causal, with effect sizes assigned to observed genotypes. This design choice is essential for the controlled factorial analysis that is central to our study: it ensures that the conditional distribution shift (controlled by ρ) and the marginal distribution shift (determined by allele frequency differences) are independently manipulable parameters, free from the confounding that would arise if prediction accuracy also depended on LD-mediated tagging imprecision. This separation enables clean attribution of performance differences to specific types of distribution shift, which is the primary objective of our experimental design. The degree of data representation bias is controlled by the proportion of the DDP (λ) in a synthetic dataset and the conditional distribution shift is controlled by the genetic correlation between the ancestry groups (ρ).

The marginal distribution shift is determined by the allele frequency differences of the genetic variants across ancestry groups, which can be measured by the genetic distance defined as *d_ij_* = 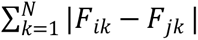, where 𝐹_𝑖k_ is the frequency of the minor allele of k^𝑡ℎ^ SNP in the ancestry group *i*, and 𝐹_𝑗k_is the frequency of the same allele in the ancestry group *j*. This metric captures the cumulative differences in allele frequencies between the two groups. We also used the maximum mean discrepancy (MMD), a statistical metric for quantifying the distance between two probability distributions [57], to measure the marginal distribution shift between ancestry groups.

### Machine learning methods

Machine learning experiments were conducted using a neural network model implemented with Keras, employing a specific configuration of layers and hyperparameters to optimize training. The input layer consisting of N nodes (N is the number of input genotype features) is connected to a hidden layer of 200 nodes with ReLU activation. This hidden layer is followed by layer normalization, which stabilizes and accelerates training by normalizing activations across features, and a Dropout layer with a 10% dropout rate, reducing overfitting by randomly setting a proportion of neuron activations to zero during training [58]. The model’s output layer uses linear activation for regression and sigmoid activation for classification. The model’s configuration includes regularization penalties (L1 and L2), each set at 0.001 to impose constraints on the model coefficients, preventing large weights and promoting sparsity. The Adam optimizer, with a learning rate of 0.001, was used to compile the model. The training process incorporated several techniques to enhance performance and prevent overfitting. Early stopping was used to monitor the validation metric and stop training when it ceased to improve, with a patience of 30 epochs, restoring the best weights observed during training [59]. Additionally, the Keras ReduceLROnPlateau callback was employed to reduce the learning rate by a factor of 0.3 if the validation loss did not improve for four consecutive epochs.

Fine-tuning, a transfer learning method for adapting pre-trained models to new domains to address the discrepancies between the source and target domain [60], was used to improve the machine learning performance on DDPs. This adaptation process enhances model performance on the target domain by adjusting the neural network’s weights [60]. Additionally, fine-tuning facilitates the utilization of a pre-trained model in scenarios where the target domain contains limited data [61]. During fine-tuning, the initial learning rate for the Adam optimizer was reduced by a factor of 10. Using a lower learning rate during fine-tuning enables smaller, incremental updates to the pre-trained model parameters, allowing the model to smoothly optimize the weights learned from the source domain for the target domain. This prevents drastic updates to previously acquired weights, thereby preserving and effectively transferring the representation already learned from the source domain, enhancing prediction performance for the target domain with limited data.

### Model performance evaluation

Each dataset (consisting of EUR and a DDP) was partitioned with a stratified train-test split. Individuals were randomly assigned to the training (75 %) and testing (25 %) subsets within each ancestry stratum, ensuring that both training and testing subsets preserved the same λ value. The median R^2^ and AUROC values from 40 independent runs were used as the performance metrics for the continuous and binary phenotype prediction tasks, respectively. For continuous phenotype prediction, the performance of mixture learning for EUR and DDP was Mix1= median(R_Mix_EUR__^2^) and Mix2= median(R_Mix_DDP__^2^), the performance of independent learning for EUR and DDP was Ind1=median(R_Ind_EUR__^2^) and Ind2=median(R_Ind_DDP__^2^), and the performance of transfer learning was TL=median(R^2^_TL_). Similarly, for binary phenotype prediction, the performance of mixture learning for EUR and the DDP was Mix1 = median(AUROC_Mix_EUR__) and Mix2= median(AUROC_Mix_DDP__), the performance of independent learning for the EUR and DDP was Ind1=median(AUROC_Ind_EUR__) and Ind2= median(AUROC_Ind_DDP__), and performance of transfer learning was TL=median(AUROC_TL_).

## Results

### Multi-population machine learning experiments on genotype-phenotype datasets

We conducted machine learning experiments on synthetic genotype-phenotype datasets representing five continental ancestries (AFR, AMR, EAS, EUR, and SAS) with controlled representation bias, as well as marginal and conditional distribution shifts between the EUR and the other ancestry groups (See Methods). The machine learning tasks are to predict phenotype values from genotype data using datasets consisting of two ancestry groups: EUR and a non-EUR group of smaller sample size, which represents the data-disadvantaged population (DDP). Each dataset contains 100,000 individuals and 3,000 genetic variants.

### Parameter space representing data representation bias and conditional distribution shift

To study how data representation bias and conditional distribution shift affect multi-population machine learning, we mapped the performance landscapes of mixture learning, independent learning, and transfer learning onto a two-dimensional space defined by parameters λ and ρ (**Fig. 1**). Here, λ is the proportion of the DDP in the data and ρ is the correlation of effect sizes of the genetic variants between ancestry groups, which are inversely related to data representation bias and conditional distribution shift, respectively. A synthetic dataset was generated at each point on a grid of the two parameters: λ = 0.1, 0.2, 0.3, 0.4, and 0.5, and ρ = 0, 0.2, 0.4, 0.6, 0.8, and 1. This parameter grid essentially covers the full ranges of data representation bias, from extreme data representation bias (90% EUR and 10% DDP) to equal representation (50% EUR and 50% DDP), and spans the entire range of conditional distribution shift, from completely uncorrelated to identical effect sizes between the ancestry groups. We conducted machine learning experiments on these synthetic datasets to systematically explore how data representation bias and conditional distribution shift influence performance disparities and their mitigation.

**Fig. 1.**
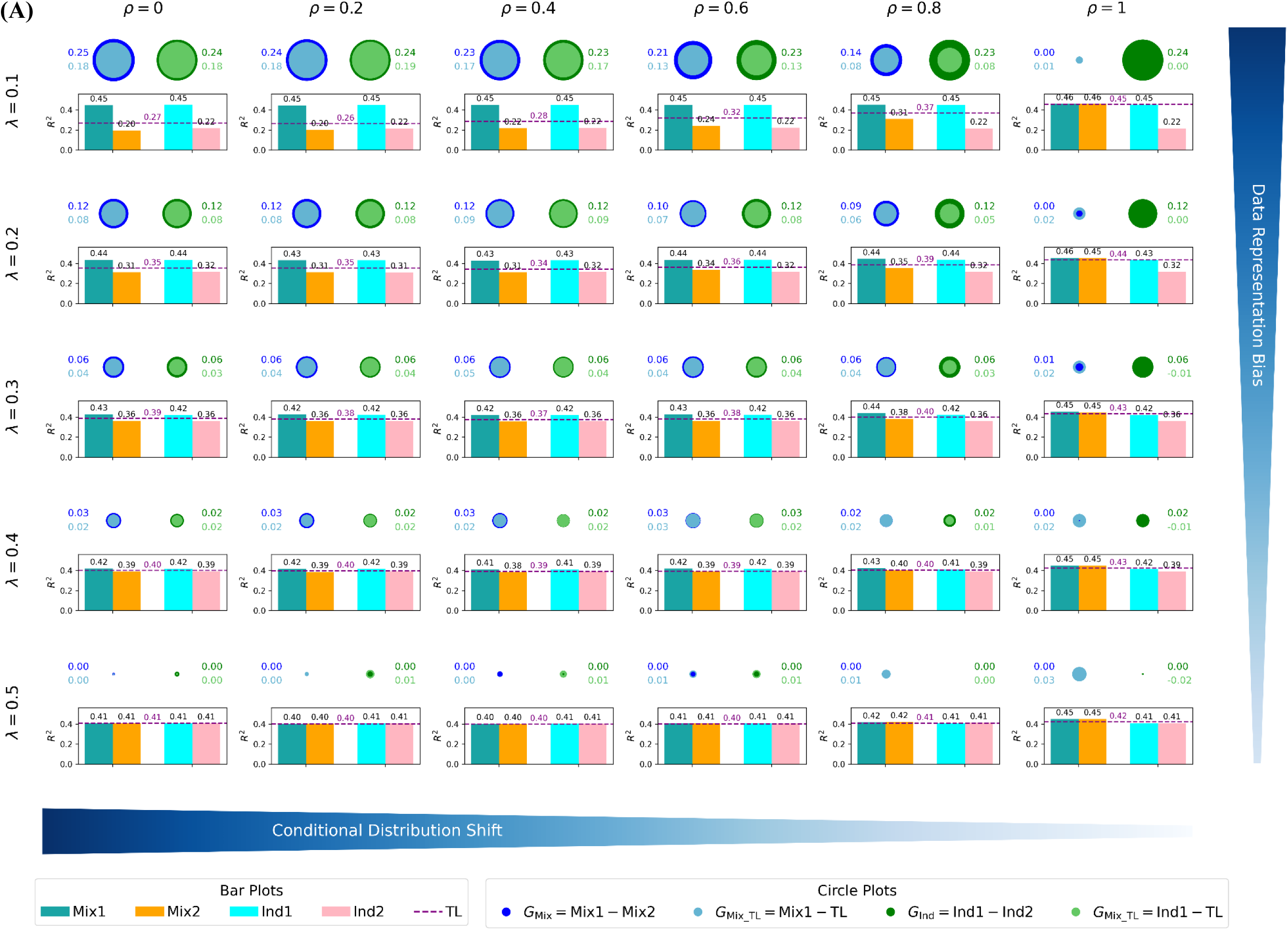
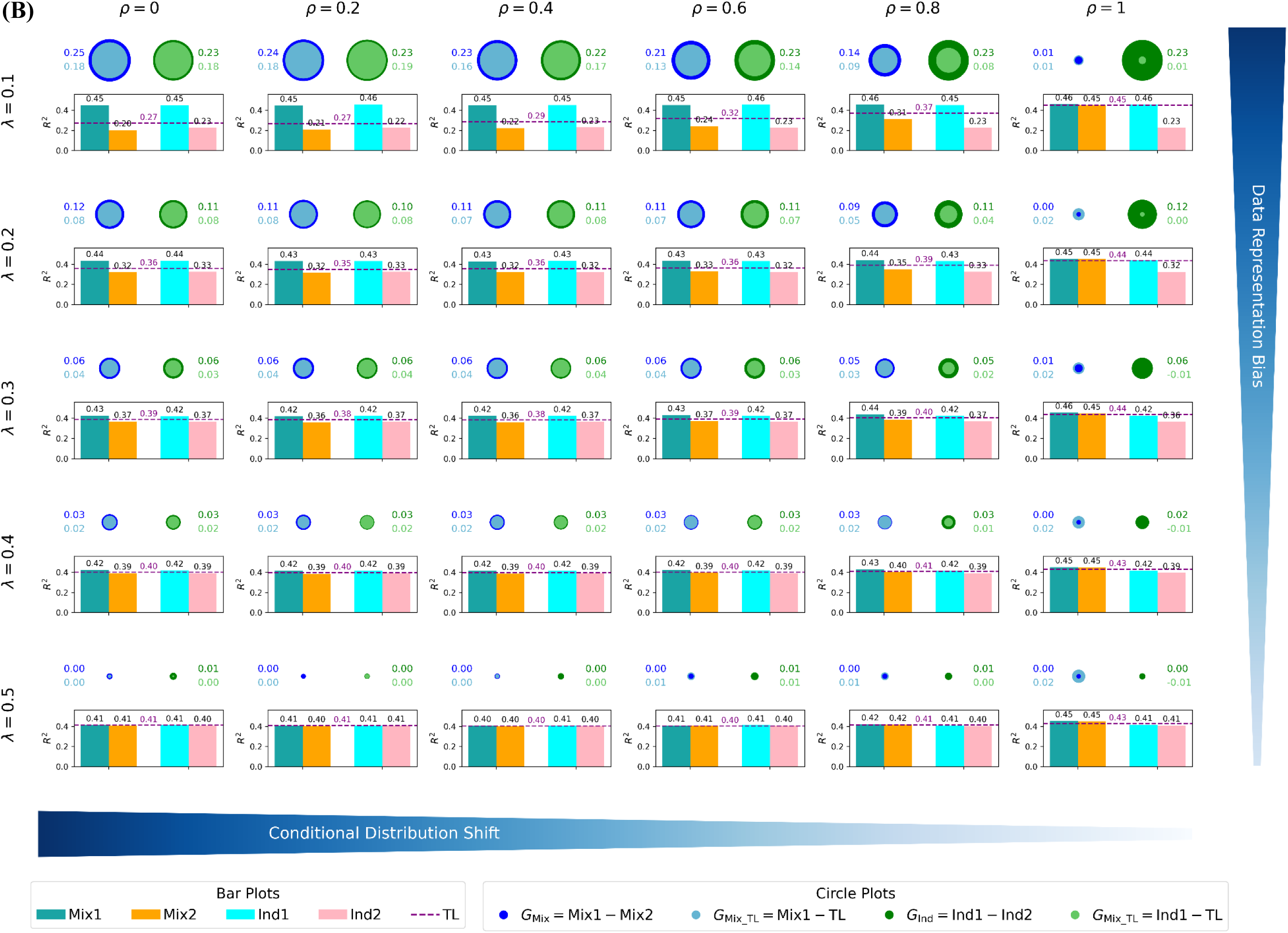
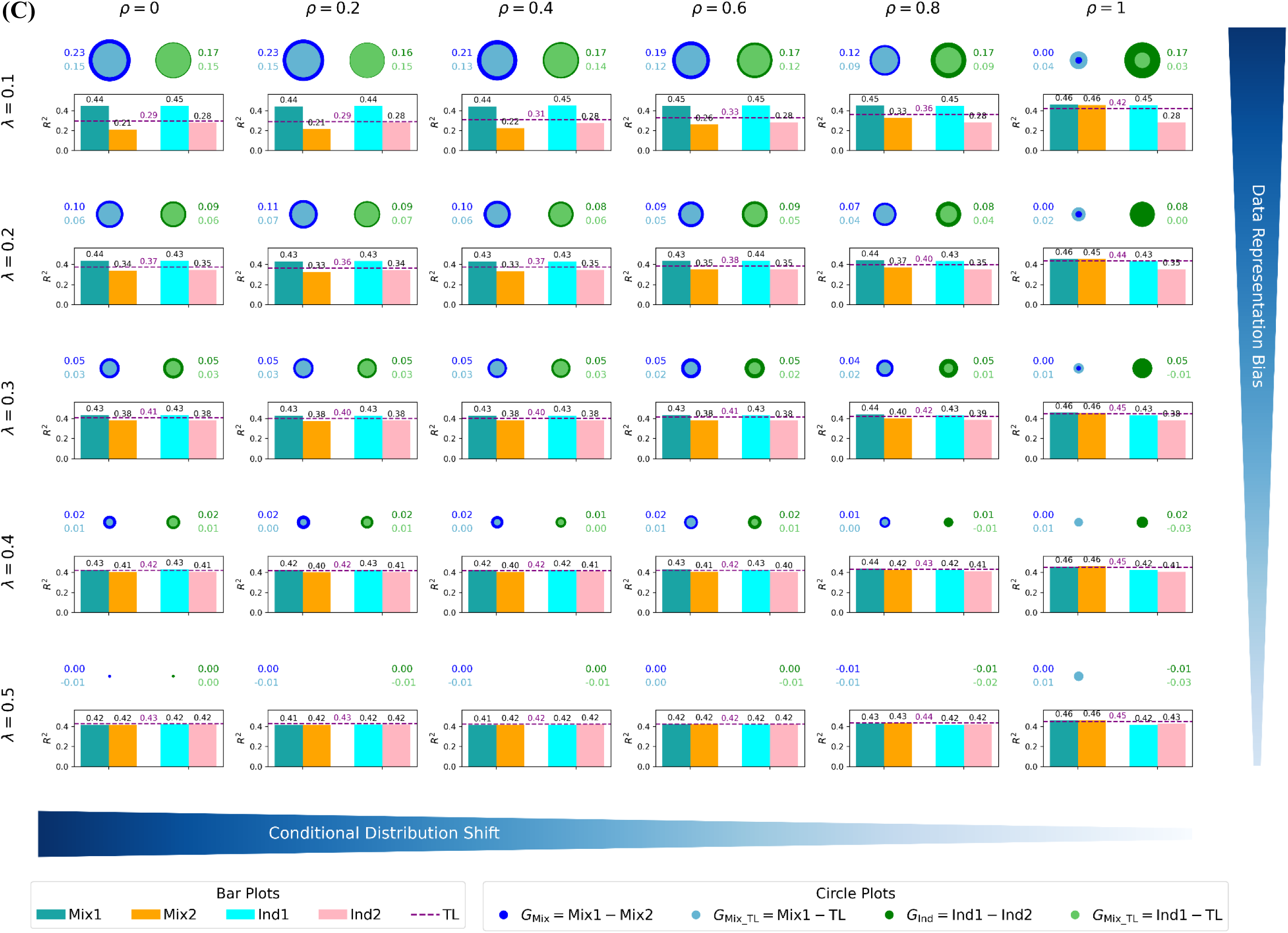
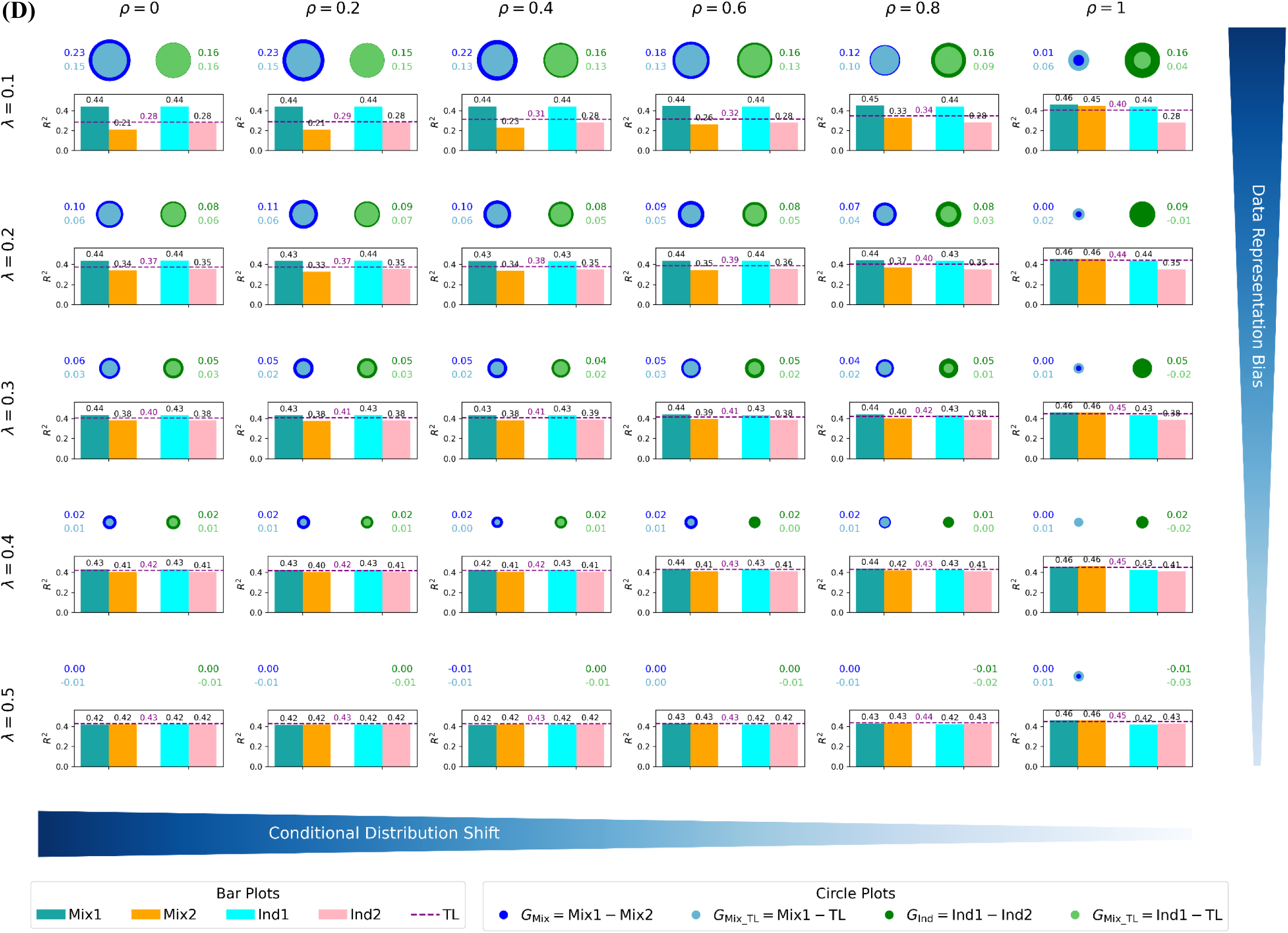
Mapping multi-ancestry machine learning model performance landscapes for the populations consisting of **(A)** EUR and AMR, **(B)** EUR and SAS, **(C)** EUR and EAS, and **(D)** EUR and AFR (*continuous* phenotype, h^2^=0.5, number of SNPs=3,000, total population size=100,000). The hierarchical grid of subplots illustrates the variation in machine learning model performance across a parameter space defined by data representation bias and conditional distribution shift. In each subplot, the bar charts and dashed line show performance of different multi-ancestry machine learning approaches. Mix1 and Mix2 represent the performance of mixture learning for EUR and DDP, Ind1 and Ind2 represent the performance of independent learning for EUR and DDP, and the dashed line represents the performance of transfer learning. The blue and light blue concentric circles represent G_Mix_ and G_Mix_TL_, and the green and light green concentric circles represent G_Ind_ and G_Ind_TL_.

### Impacts of data representation bias and distribution shifts

For the EUR population, the performance of both the mixture and independent learning models remains largely stable, with variations of less than 10% across the parameter space (see Mix1 and Ind1, the first and third bars in each panel of **Fig. 1**). This stability likely reflects that the EUR models were trained on at least 50% of the data (or 50,000 individuals), approaching data sufficiency (“data saturation”), beyond which additional data does not improve the model’s predictive capability significantly [62, 63].

For the DDP, the performance of the mixture learning models (Mix2, the second bar in each panel of **Fig. 1**) is a function of data representation bias and conditional distribution shift, increasing as either factor decreases; the performance of the independent learning models (Ind2, the fourth bar in each panel of **Fig. 1**) also increases as data representation bias decreases but is unaffected by the conditional distribution shift.

The disparity gap from mixture learning is defined as G_Mix_ = Mix1 − Mix2, and the disparity gap from independent learning is defined as G_Ind_ = Ind1 − Ind2 . G_Mix_ decreases as data representation bias reduces (blue circles in **Fig. 1**), approaching 0 when there is no data representation bias (λ = 0.5) or conditional distribution shift (ρ = 1). G_Ind_ also decreases as data representation bias reduces but remains largely unchanged across different levels of conditional distribution shift (green circles in **Fig. 1**).

The performance of transfer learning is a function of data representation bias and conditional distribution shift, increasing as either factor decreases (see TL, the purple line in each panel of **Fig. 1**). When using transfer learning performance for the DDP, the disparity gaps are: G_MixTL_ = Mix1 − TL and G_IndTL_ = Ind1 − TL (light blue and light green circles in **Fig. 1**). For the DDP, transfer learning outperforms both mixture (Mix2) and independent learning (Ind2) in most regions of the parameter space, thereby reducing the disparity gaps. In these regions, G_MixTL_ < G_Mix_ and G_IndTL_ < G_Ind_, demonstrating that transfer learning is effective for disparity mitigation.

We define disparity regions in the λ-ρ space as those where the disparity gap is at least 0.01. For mixture learning, the disparity region spans ρ from 0 to 0.8 and λ from 0.1 to 0.4. Transfer learning reduces disparity gaps across the entire disparity region (**Fig. 1**), with the larger reductions occurring in areas of high representation bias and large conditional distribution shifts. For independent learning, the disparity region is broader in the ρ dimension, extending from 0 to 1, while remaining between 0.1 and 0.4 in the λ dimension. Transfer learning also reduces disparity gaps across the entire disparity region, with larger reductions observed in areas of high data representation bias but small conditional distribution shifts (**Fig. 1**).

We then investigated how marginal distribution shift affects multi-population machine learning. The marginal distribution shift arises from allele frequency differences between ancestry groups. We computed genetic distance and maximum mean discrepancy (MMD) as measures of marginal distribution shift (see Methods). The normalized matrix integrating genetic distance and MMD is visualized in **Fig. 2**. As expected, both measures indicate that the marginal distribution shift of each DDP relative to EUR increases progressively from AMR, SAS, EAS, to AFR. To assess whether the observed performance landscape in the λ-ρ space varies with marginal distribution shift, we conducted the same machine learning experiments using four combinations of EUR and DDPs: EUR+ AMR, EUR+ SAS, EUR+EAS, and EUR+ AFR. Across all EUR+DDP combinations, performance patterns with respect to data representation bias and conditional distribution shift are highly consistent (**Fig. 1, A-D**), indicating that the effect of marginal distribution shift on multi-population machine learning performance is limited.

**Fig. 2.**
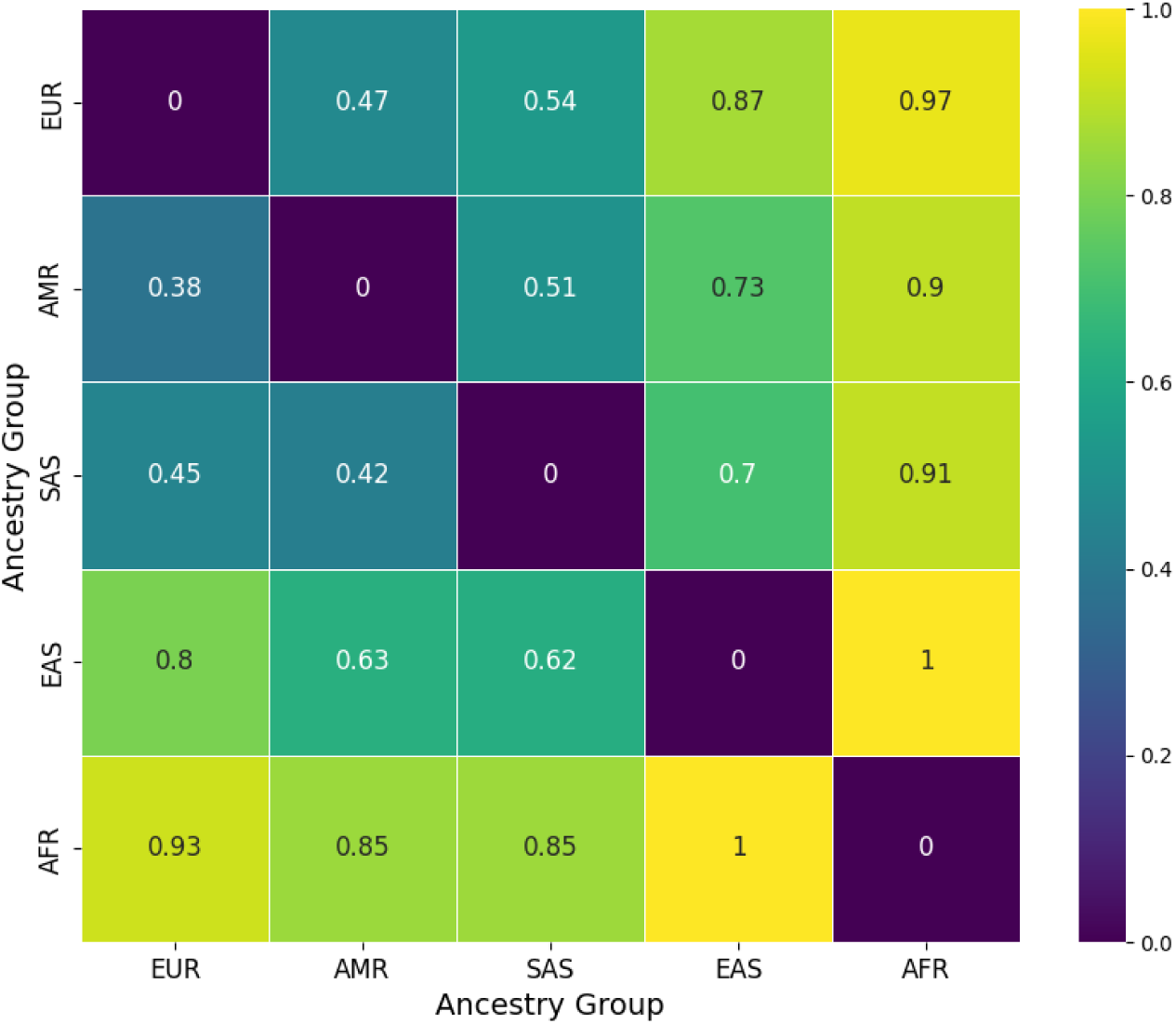
Normalized genetic distance (upper triangle) and marginal distribution shift measured by the maximum mean discrepancy (MMD) values (lower triangle) between ancestry groups.

In summary, our experiments show how multi-population machine learning performance varies with data representation bias and distribution shifts:

- **Joint effects of data representation bias and conditional distribution shifts.** In mixture learning, DDP performance improves when either data representation bias or conditional distribution shift is reduced. In independent learning, performance for the DDP improves only with reduced data representation bias and is unaffected by conditional distribution shift. Consequently, the mixture learning disparity gap decreases when either factor is reduced, whereas the independent learning disparity gap decreases only with reduced data representation bias but remains largely unchanged across different levels of conditional distribution shifts. Transfer learning generally outperforms mixture and independent learning for the DDP, thereby reducing disparity gaps.
- **Limited influence of marginal distribution shift.** Experiments across four EUR+DDP combinations show that although these pairs of ancestry groups have different marginal distribution shifts, the overall patterns of model performance with respect to data representation bias and conditional distribution shift remain consistent. Thus, marginal distribution shift does not substantially affect multi-population machine learning performance.

We further tested the robustness of these findings using datasets generated at different heritability levels (h^2^=0.5, **Fig. 1**; and h^2^=0.25, **Fig. S1**), as well as in a different experimental setting: binary-phenotype prediction with a smaller dataset consisting of 500 SNPs and 10,000 individuals (with a case-to-control ratio of 1:4). These findings remained consistent across all settings, as shown by the 16 λ-ρ space performance landscape atlases in **Fig. 1**, **Fig. 3, Fig. S1** and **Fig. S2**. To quantify variation in multi-population performance across diverse conditions, we assessed similarities among the 16 performance landscapes. For each λ-ρ space performance landscape atlas, we constructed a single vector by concatenating the performance metrics [Mix1, Mix2, Ind1, Ind2, and TL] across all λ-ρ grid points, then computed Pearson and Spearman correlations between these atlas-level vectors. The resulting correlation matrix (**Table 1**) indicates high overall consistency, with average Pearson and Spearman correlation coefficients of 0.940 and 0.907, respectively, across different ancestry-group pairs, heritability levels, and machine-learning tasks (phenotype type and dataset size), thereby validating the robustness of the λ-ρ space performance landscape underlying our conclusions.

**Fig. 3.**
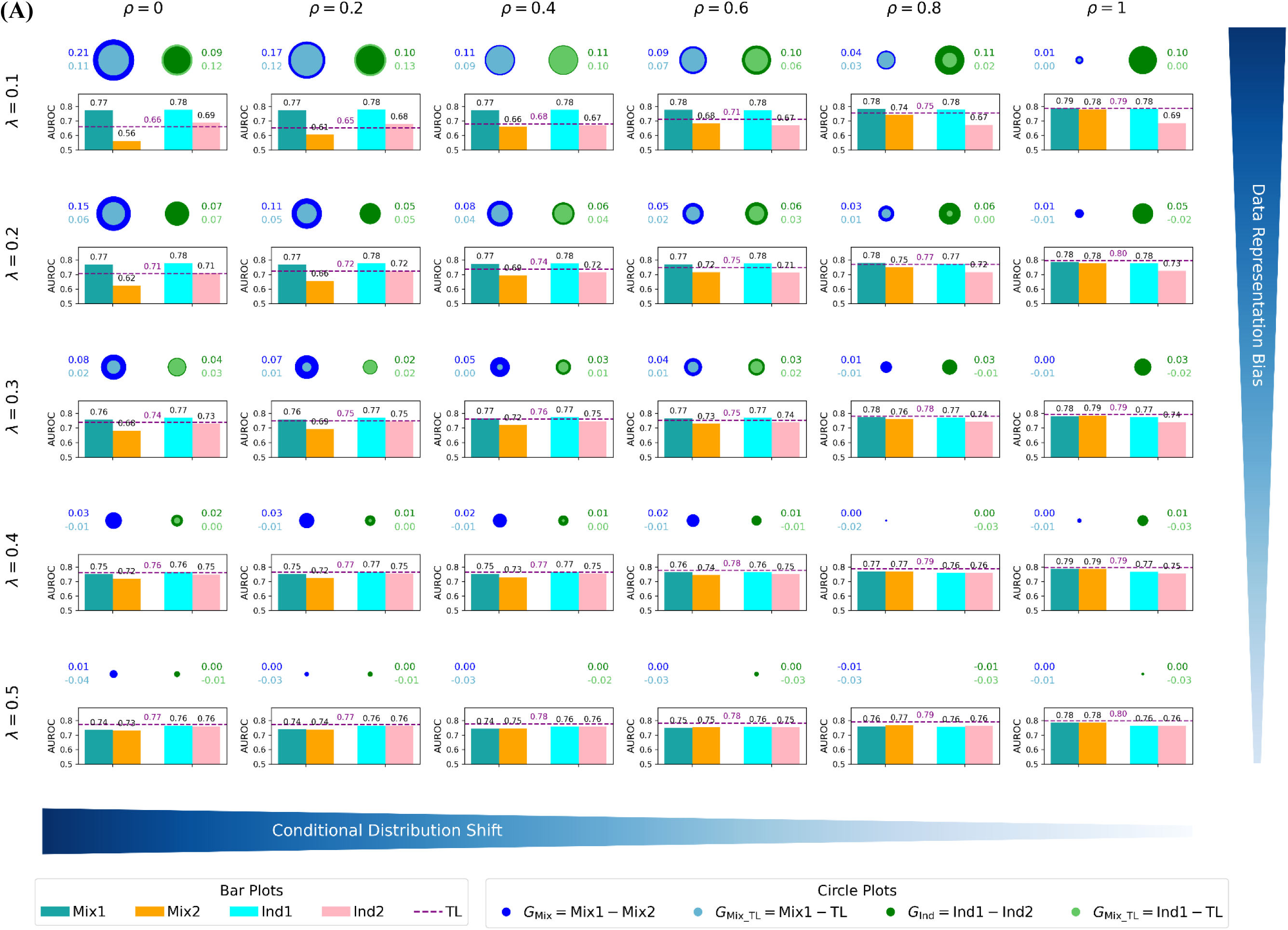
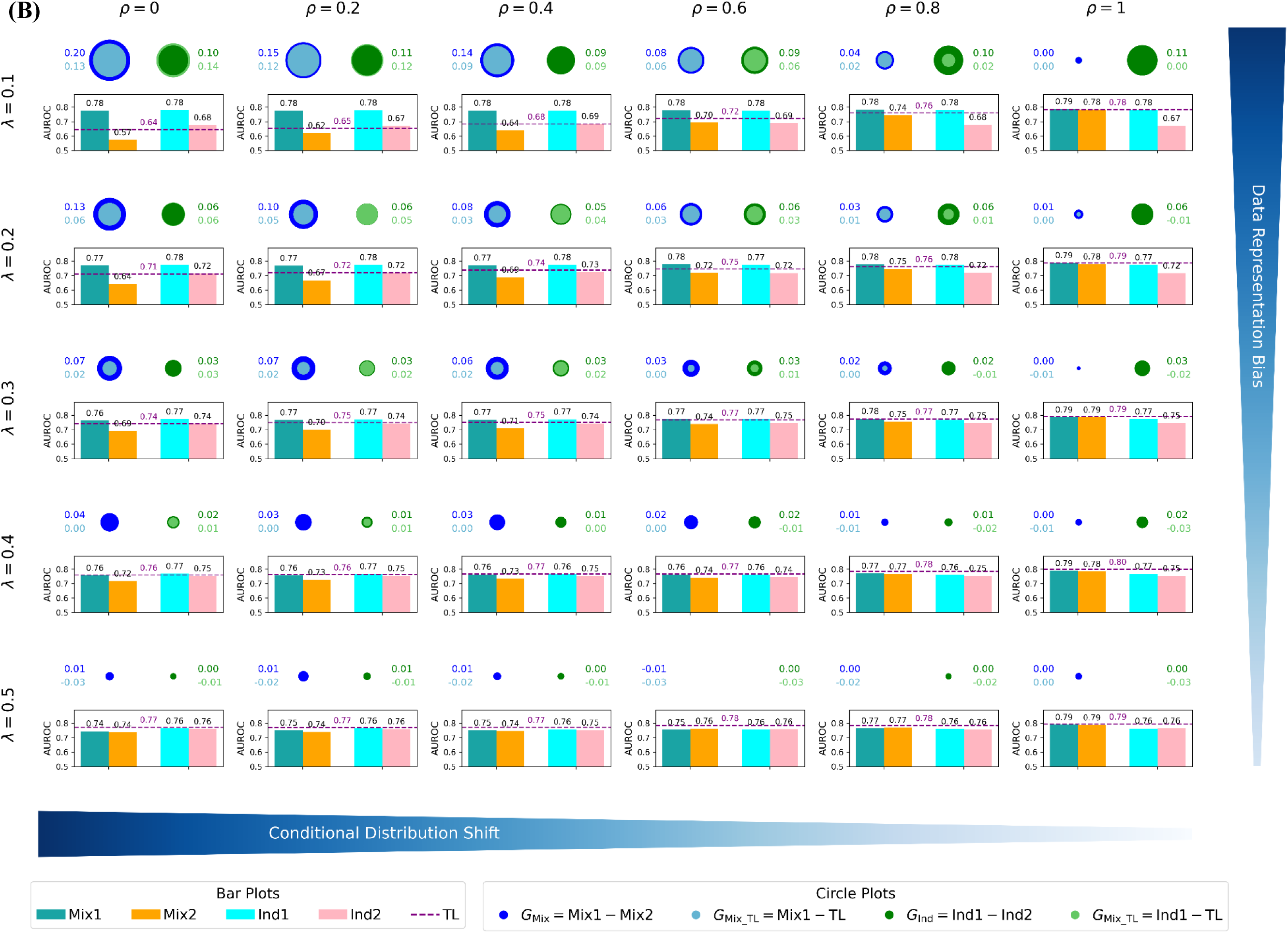
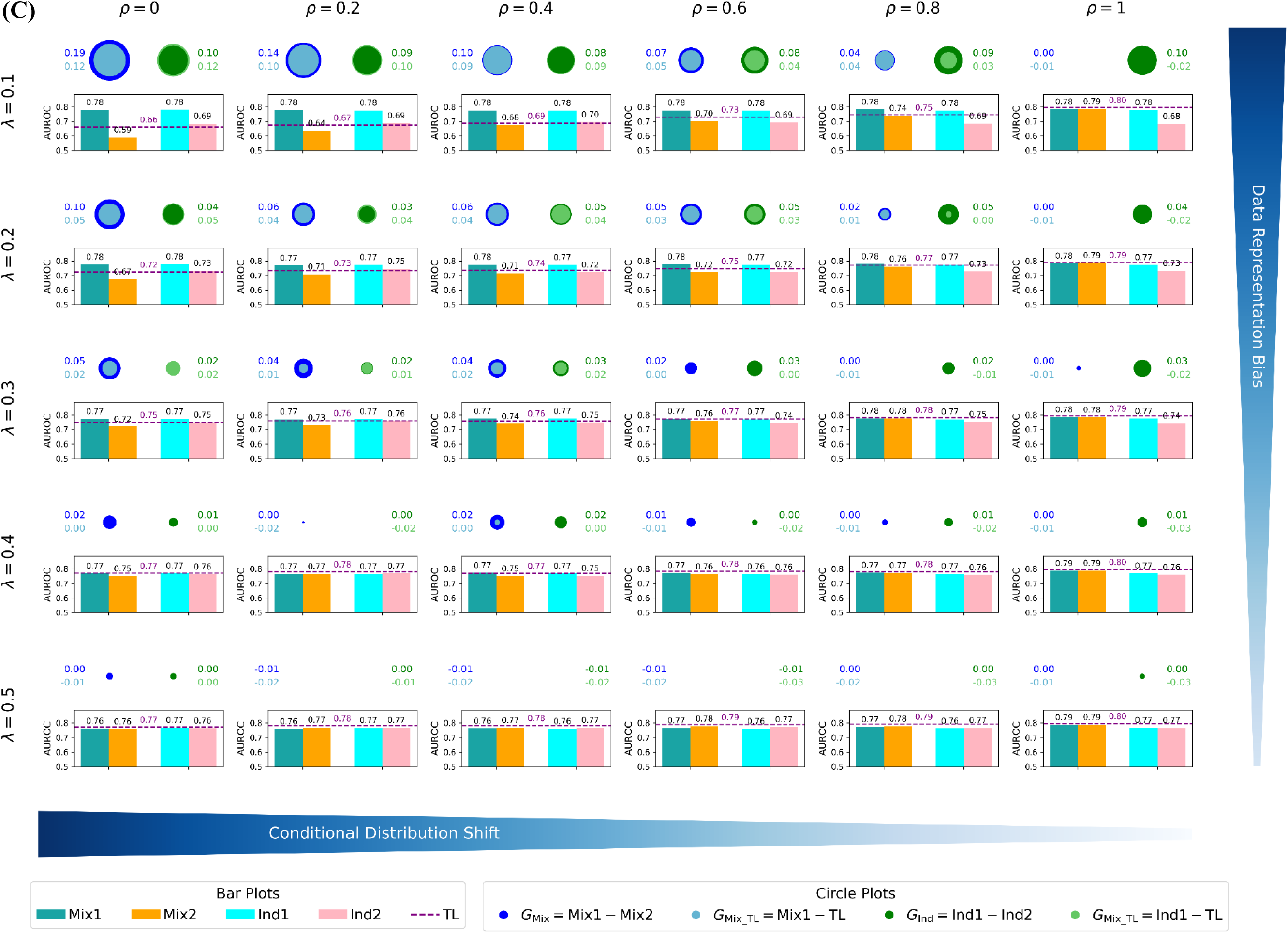
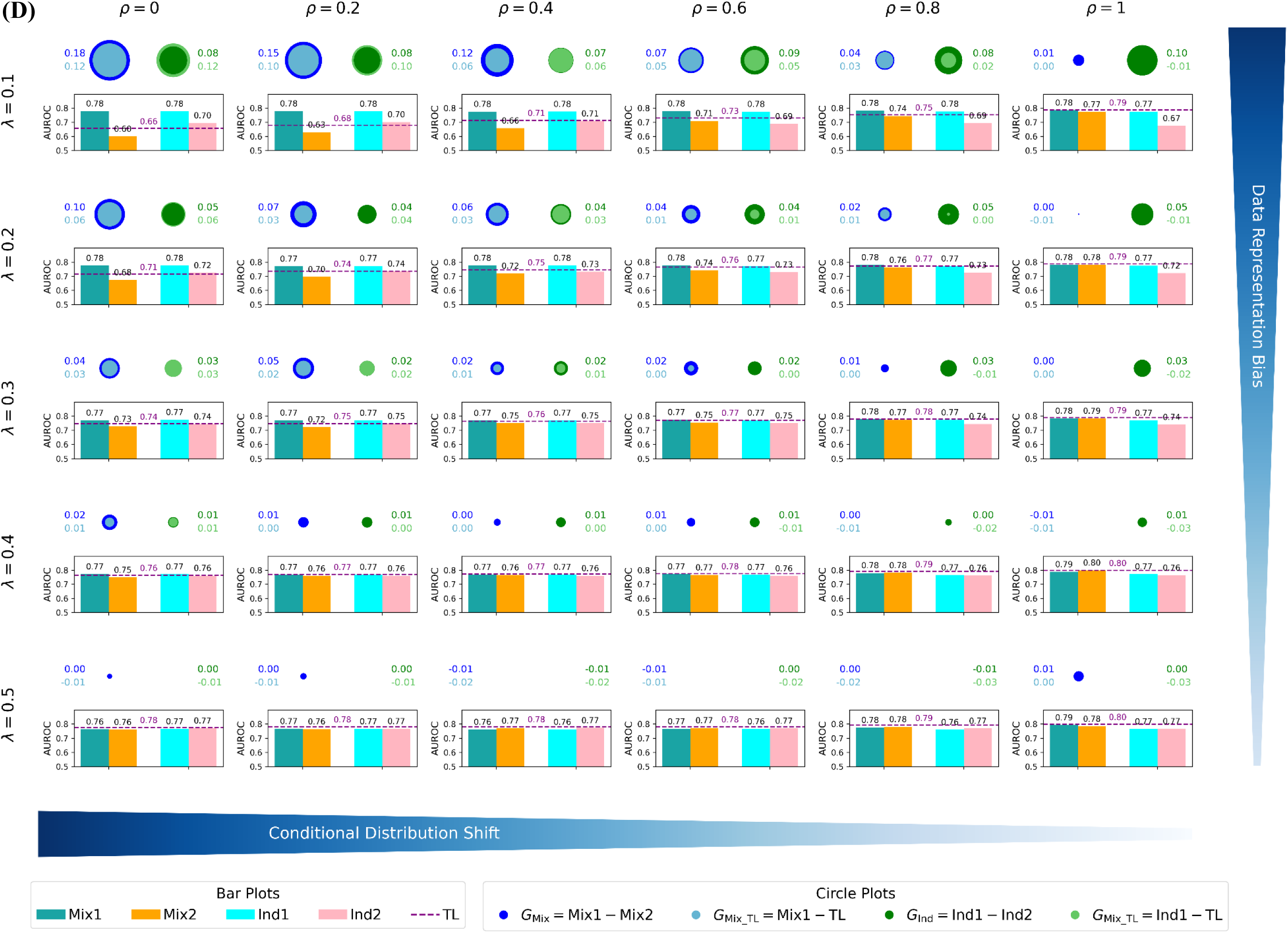
Mapping multi-ancestry machine learning model performance landscapes for the populations consisting of **(A)** EUR and AMR, **(B)** EUR and SAS, **(C)** EUR and EAS, and **(D)** EUR and AFR (*binary* phenotype, h^2^=0.5, number of SNPs=500, total population size=10,000). The hierarchical grid of subplots illustrates the variation in machine learning model performance across a parameter space defined by data representation bias and conditional distribution shift. In each subplot, the bar charts and dashed line show performance of different multi-ancestry machine learning approaches. Mix1 and Mix2 represent the performance of mixture learning for EUR and DDP, Ind1 and Ind2 represent the performance of independent learning for EUR and DDP, and the dashed line represents the performance of transfer learning. The blue and light blue concentric circles represent G_Mix_ and G_Mix_TL_, and the green and light green concentric circles represent G_Ind_ and G_Ind_TL_.

**Table 1.** Correlation of multi-population machine learning performance across different conditions.

|  |  |  | EUR+AFR |  |  |  | EUR+AMR |  |  |  | EUR+EAS |  |  |  | EUR+SAS |  |  |  |
| --- | --- | --- | --- | --- | --- | --- | --- | --- | --- | --- | --- | --- | --- | --- | --- | --- | --- | --- |
| | | | $h^2=0.25$ | | $h^2=0.5$ | | $h^2=0.25$ | | $h^2=0.5$ | | $h^2=0.25$ | | $h^2=0.5$ | | $h^2=0.25$ | | $h^2=0.5$ | |
|  |  |  | CP | BP | CP | BP | CP | BP | CP | BP | CP | BP | CP | BP | CP | BP | CP | BP |
| EUR+AFR | $h^2=0.25$ | CP | 1.000 | 0.934 | 0.985 | 0.961 | 0.978 | 0.906 | 0.946 | 0.934 | 0.996 | 0.916 | 0.981 | 0.949 | 0.980 | 0.905 | 0.950 | 0.944 |
|  |  | BP | 0.847 | 1.000 | 0.915 | 0.969 | 0.901 | 0.938 | 0.873 | 0.959 | 0.921 | 0.970 | 0.914 | 0.969 | 0.903 | 0.946 | 0.878 | 0.965 |
| | $h^2=0.5$ | CP | 0.976 | 0.869 | 1.000 | 0.948 | 0.986 | 0.871 | 0.972 | 0.902 | 0.985 | 0.901 | 0.997 | 0.935 | 0.986 | 0.873 | 0.977 | 0.918 |
|  |  | BP | 0.900 | 0.936 | 0.896 | 1.000 | 0.943 | 0.937 | 0.918 | 0.962 | 0.951 | 0.961 | 0.949 | 0.988 | 0.945 | 0.944 | 0.921 | 0.974 |
| EUR+AMR | $h^2=0.25$ | CP | 0.959 | 0.826 | 0.960 | 0.873 | 1.000 | 0.871 | 0.979 | 0.898 | 0.982 | 0.904 | 0.987 | 0.930 | 0.997 | 0.871 | 0.980 | 0.915 |
|  |  | BP | 0.895 | 0.889 | 0.882 | 0.925 | 0.852 | 1.000 | 0.847 | 0.974 | 0.893 | 0.946 | 0.874 | 0.941 | 0.879 | 0.976 | 0.851 | 0.976 |
| | $h^2=0.5$ | CP | 0.924 | 0.808 | 0.950 | 0.857 | 0.950 | 0.841 | 1.000 | 0.870 | 0.950 | 0.880 | 0.981 | 0.909 | 0.978 | 0.848 | 0.998 | 0.893 |
|  |  | BP | 0.893 | 0.882 | 0.875 | 0.940 | 0.856 | 0.965 | 0.847 | 1.000 | 0.921 | 0.960 | 0.907 | 0.967 | 0.904 | 0.972 | 0.875 | 0.985 |
| EUR+EAS | $h^2=0.25$ | CP | 0.989 | 0.845 | 0.969 | 0.901 | 0.951 | 0.892 | 0.915 | 0.895 | 1.000 | 0.909 | 0.982 | 0.940 | 0.984 | 0.892 | 0.954 | 0.933 |
|  |  | BP | 0.834 | 0.950 | 0.850 | 0.937 | 0.819 | 0.909 | 0.808 | 0.908 | 0.837 | 1.000 | 0.907 | 0.966 | 0.906 | 0.945 | 0.884 | 0.961 |
| | $h^2=0.5$ | CP | 0.971 | 0.870 | 0.991 | 0.908 | 0.959 | 0.889 | 0.962 | 0.890 | 0.970 | 0.861 | 1.000 | 0.939 | 0.986 | 0.877 | 0.984 | 0.920 |
|  |  | BP | 0.899 | 0.925 | 0.892 | 0.971 | 0.867 | 0.941 | 0.861 | 0.951 | 0.900 | 0.939 | 0.908 | 1.000 | 0.933 | 0.946 | 0.911 | 0.974 |
| EUR+SAS | $h^2=0.25$ | CP | 0.958 | 0.829 | 0.958 | 0.873 | 0.991 | 0.860 | 0.949 | 0.866 | 0.953 | 0.823 | 0.957 | 0.869 | 1.000 | 0.878 | 0.980 | 0.921 |
|  |  | BP | 0.846 | 0.895 | 0.835 | 0.911 | 0.808 | 0.955 | 0.801 | 0.948 | 0.848 | 0.911 | 0.849 | 0.940 | 0.814 | 1.000 | 0.851 | 0.978 |
| | $h^2=0.5$ | CP | 0.933 | 0.817 | 0.955 | 0.864 | 0.952 | 0.853 | 0.994 | 0.859 | 0.924 | 0.815 | 0.967 | 0.869 | 0.954 | 0.812 | 1.000 | 0.897 |
|  |  | BP | 0.902 | 0.899 | 0.884 | 0.947 | 0.863 | 0.969 | 0.856 | 0.978 | 0.903 | 0.915 | 0.896 | 0.958 | 0.871 | 0.960 | 0.867 | 1.000 |

## Discussion

Data representation bias and conditional distribution shift jointly create major challenges for multi-population machine learning across various domains. When training data are skewed towards certain groups, models often perform well on those groups but poorly for underrepresented populations. This representation bias becomes especially consequential under conditional distribution shifts, i.e., when the relationship between features and outcomes differs across groups. Our findings show that conditional distribution shifts, in combination with representation bias, can significantly degrade predictive accuracy for underrepresented groups, whereas shifts in the input feature distribution alone (marginal distribution shifts) have limited impact.

This mechanism generalizes beyond genomics to domains where predictions guide testing and treatment/intervention. In dermatology imaging, for example, state-of-the-art classifiers trained on corpora dominated by lighter skin tones underperform on darker tones. Evaluation on the Diverse Dermatology Images benchmark shows significant performance drops for dark skin that narrow only after fine-tuning on a more representative dataset, indicating the impact of representation bias (insufficient dark-skin coverage) and conditional distribution shift (tone-specific presentation differences) [64]. Similarly, in chest-radiograph diagnosis, multi-dataset audits document underdiagnosis for historically underserved populations (e.g., by sex, race, socioeconomic status), with higher false-negative rates that may lead to missed disease and inappropriate care. The gaps are linked to imbalanced training data and group-specific differences in imaging and labeling, i.e., representation bias coupled with conditional distribution shift [65]. In education, nationally representative analyses show lower predictive accuracy and higher misclassification for Black and Hispanic students relative to White and Asian students. Importantly, these disparities are attributed primarily to group-specific feature-to-outcome relationships rooted in structural differences (conditional distribution shift) rather than solely to sample under-representation, which points to the need for methods that explicitly handle population-dependent mappings rather than relying on simple rebalancing [66].

Our study maps how representation bias and conditional distribution shift shape multi-population machine learning model performance, demonstrates the limited role of marginal distribution shift, and identifies conditions under which adaptation strategies yield the largest equity gains. These insights have broad implications for machine learning tasks involving population-stratified data. In general, addressing representation bias and conditional distribution differences are essential for developing fair and robust predictive models across all populations and domains. The analytical framework we employ for evaluating and mitigating disparities including systematic detection of subgroup performance gaps and algorithmic interventions such as transfer learning can be readily adopted in various contexts to enhance equitable machine learning.

## Funding

This work was supported by the US National Cancer Institute grant R01CA262296.

---

**Fig. S1.**
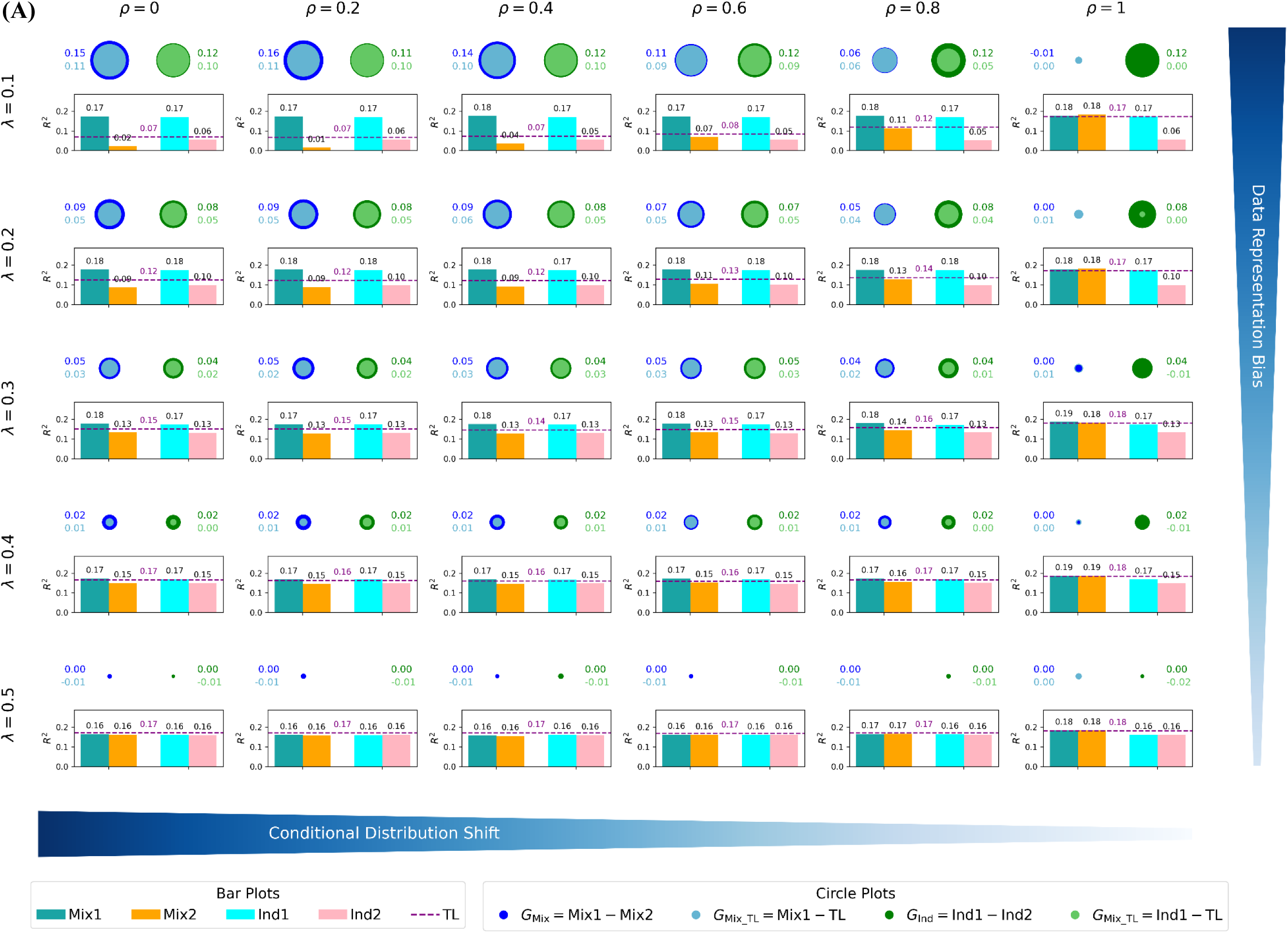
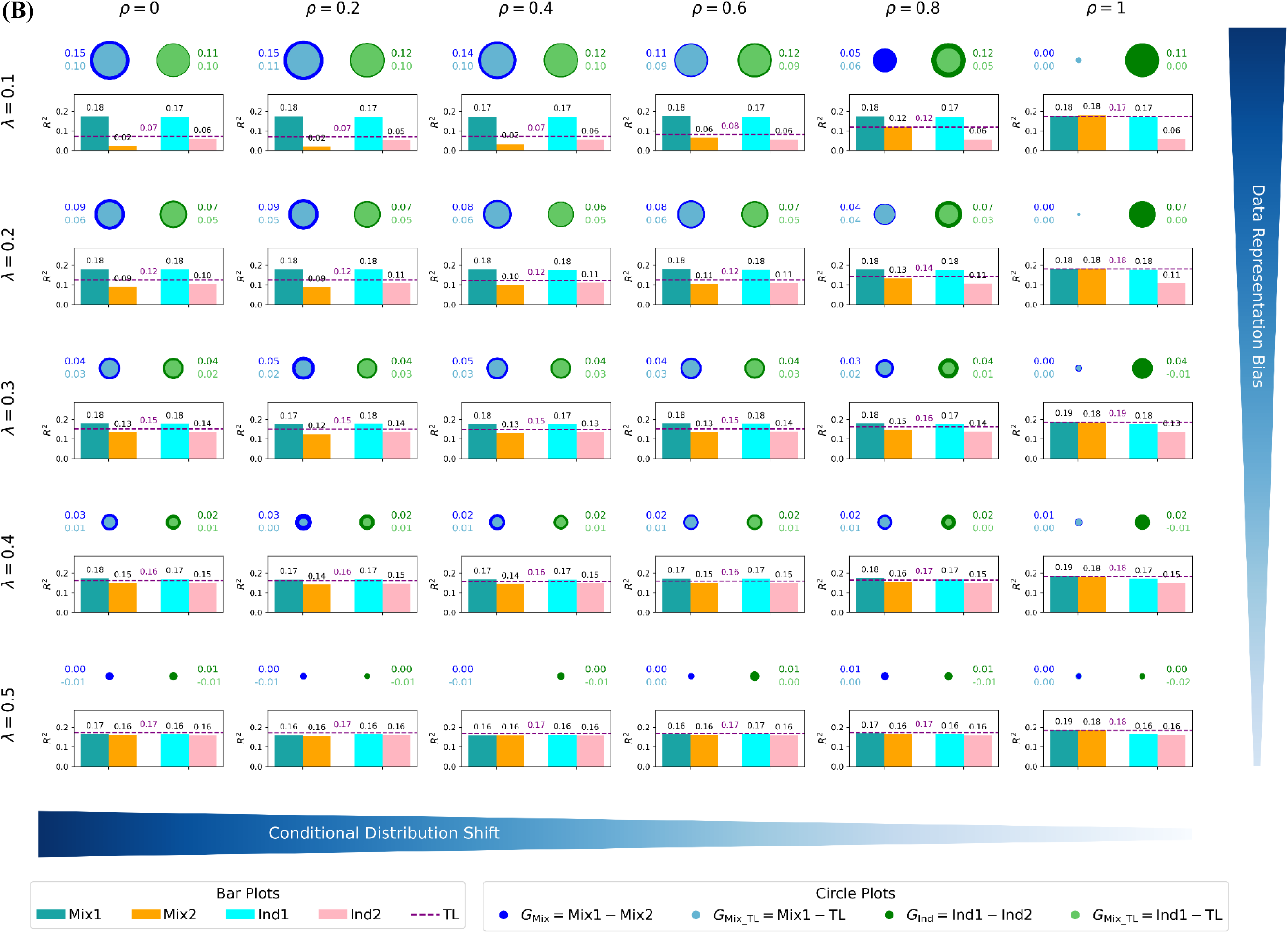
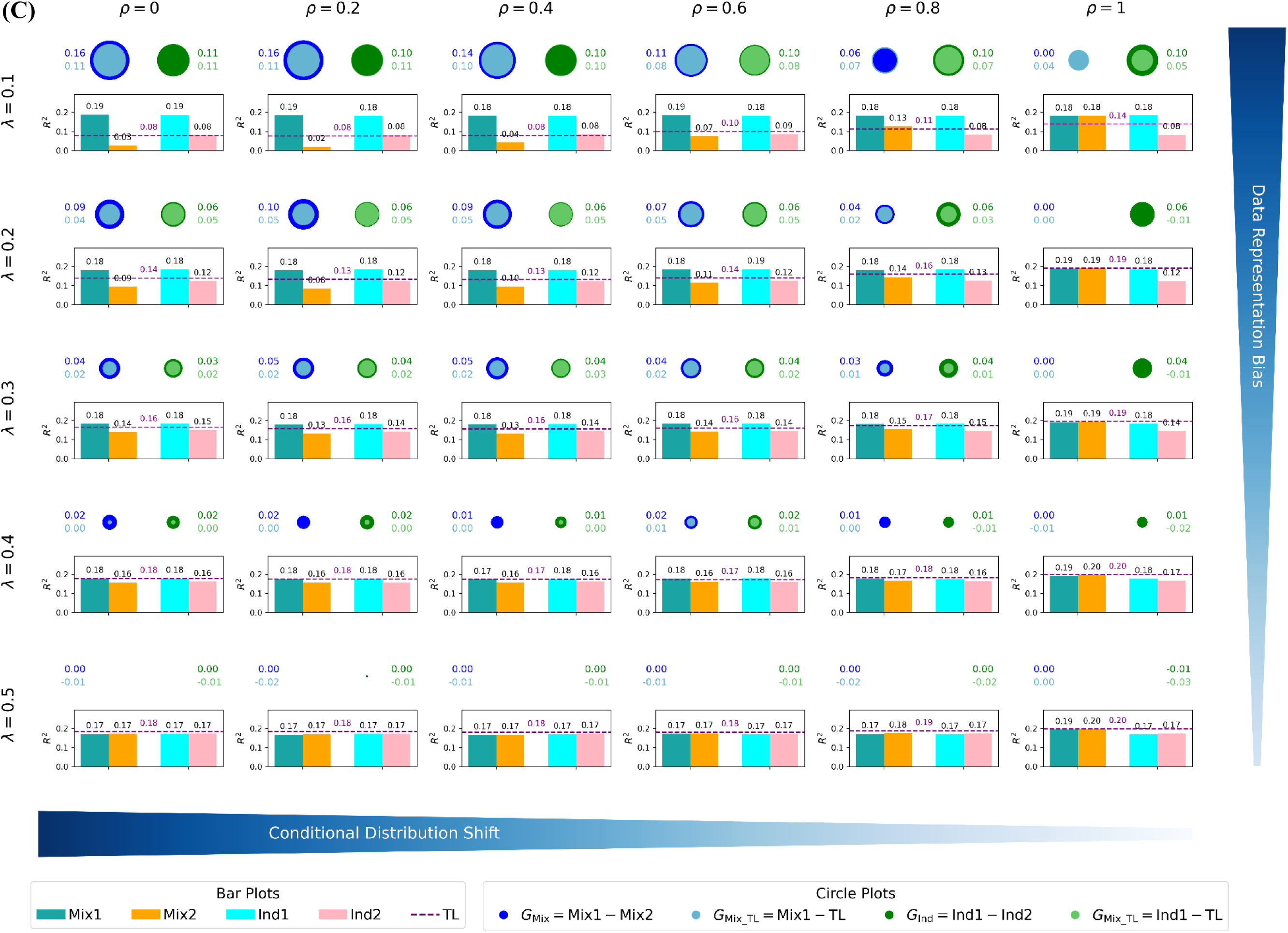
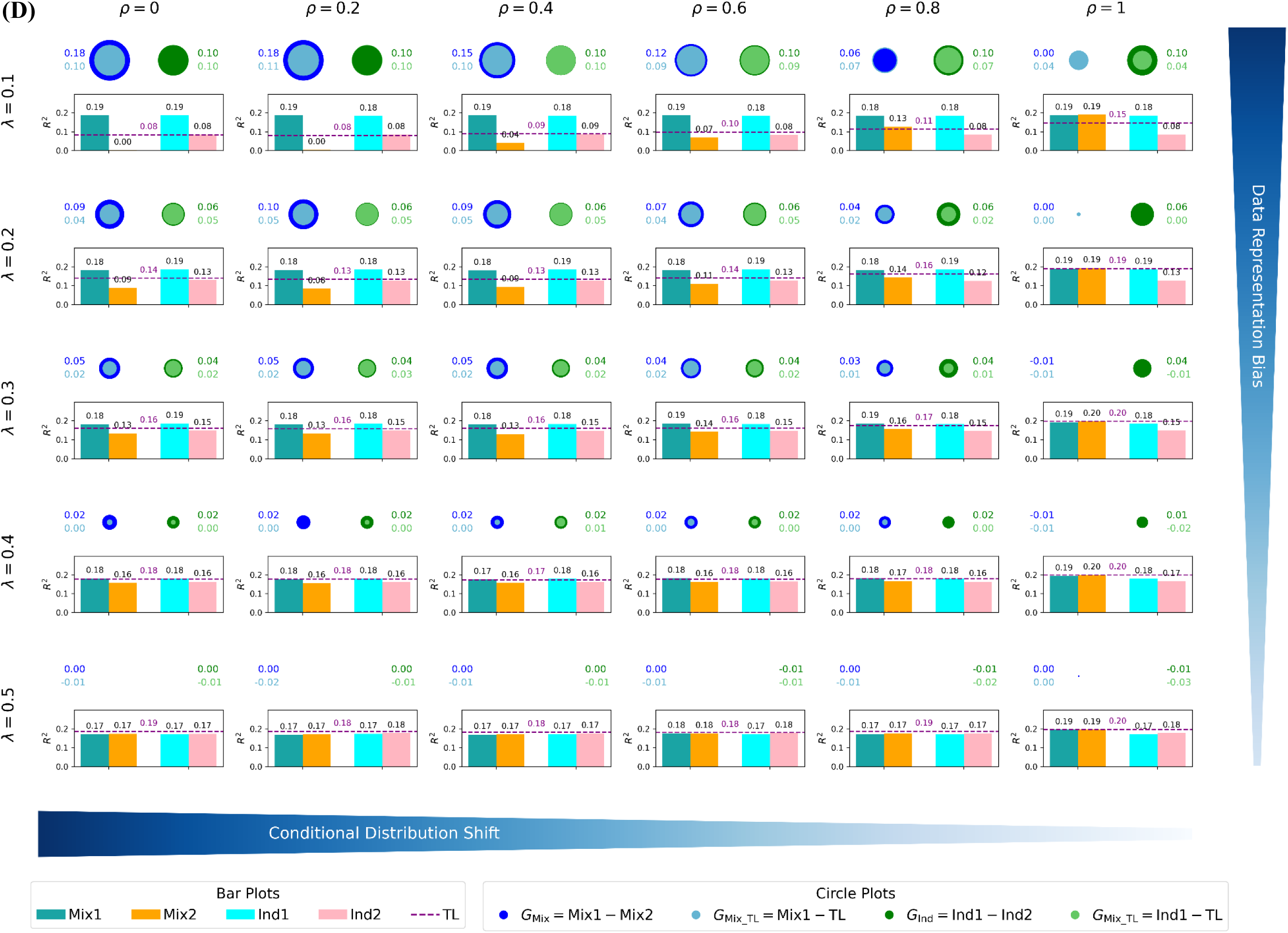
Mapping multi-ancestry machine learning model performance landscapes for the populations consisting of **(A)** EUR and AMR, **(B)** EUR and SAS, **(C)** EUR and EAS, and **(D)** EUR and AFR (*continuous* phenotype, h^2^=0.25, number of SNPs=3,000, total population size=100,000). The hierarchical grid of subplots illustrates the variation in machine learning model performance across a parameter space defined by data representation bias and conditional distribution shift. In each subplot, the bar charts and dashed line show performance of different multi-ancestry machine learning approaches. Mix1 and Mix2 represent the performance of mixture learning for EUR and DDP, Ind1 and Ind2 represent the performance of independent learning for EUR and DDP, and the dashed line represents the performance of transfer learning. The blue and light blue concentric circles represent G_Mix_ and G_Mix_TL_, and the green and light green concentric circles represent G_Ind_ and G_Ind_TL_.

**Fig. S2.**
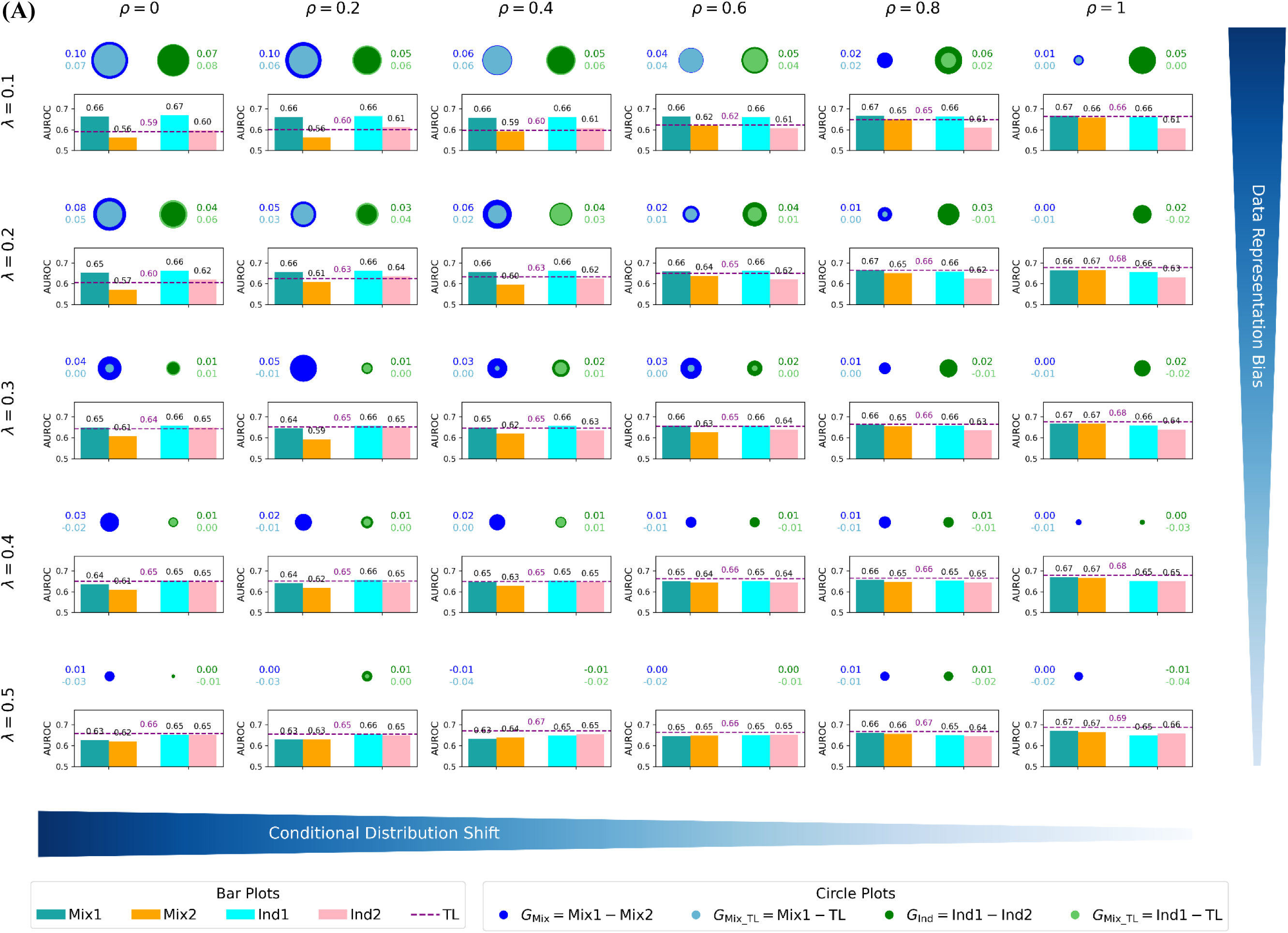
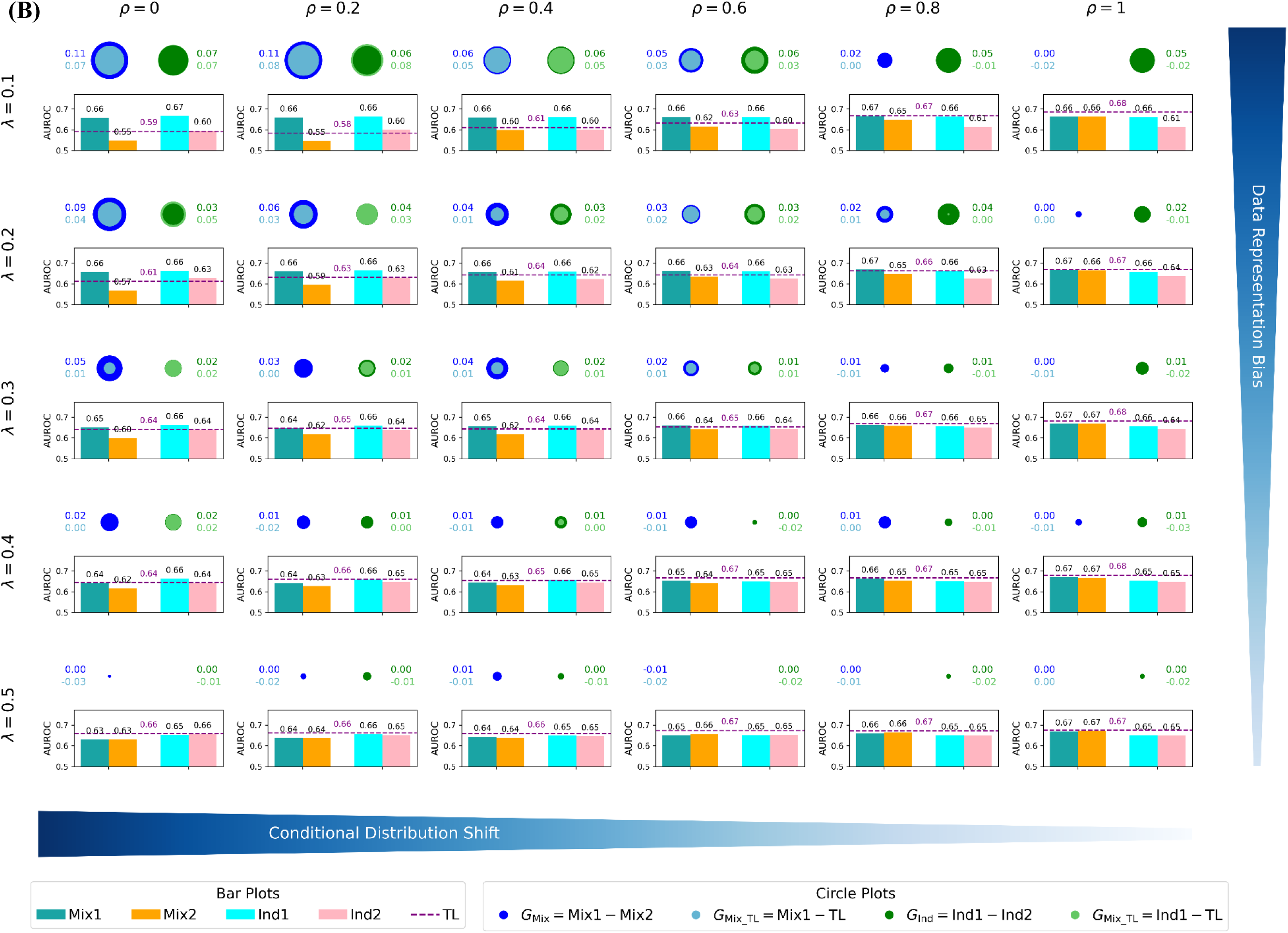
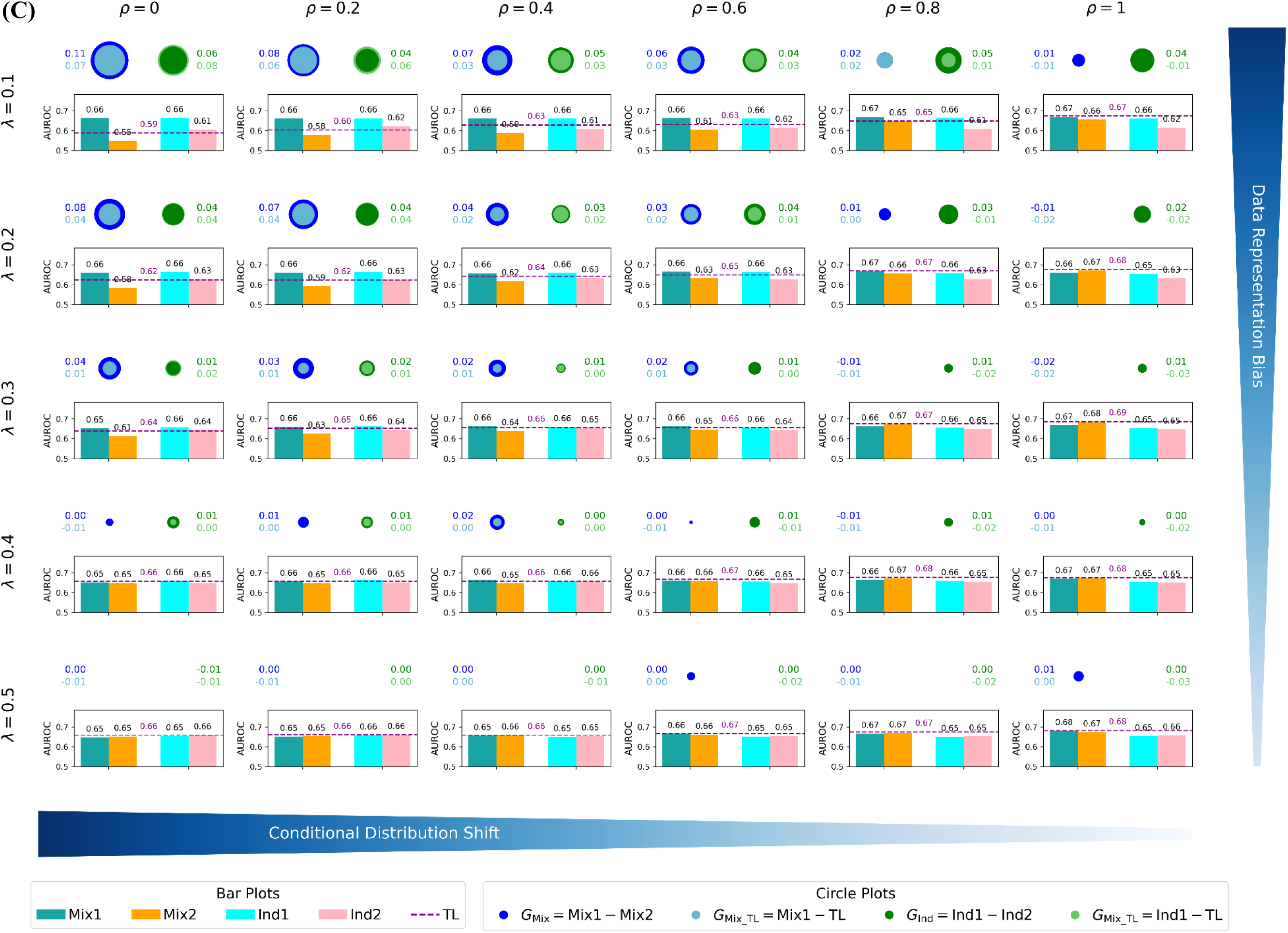
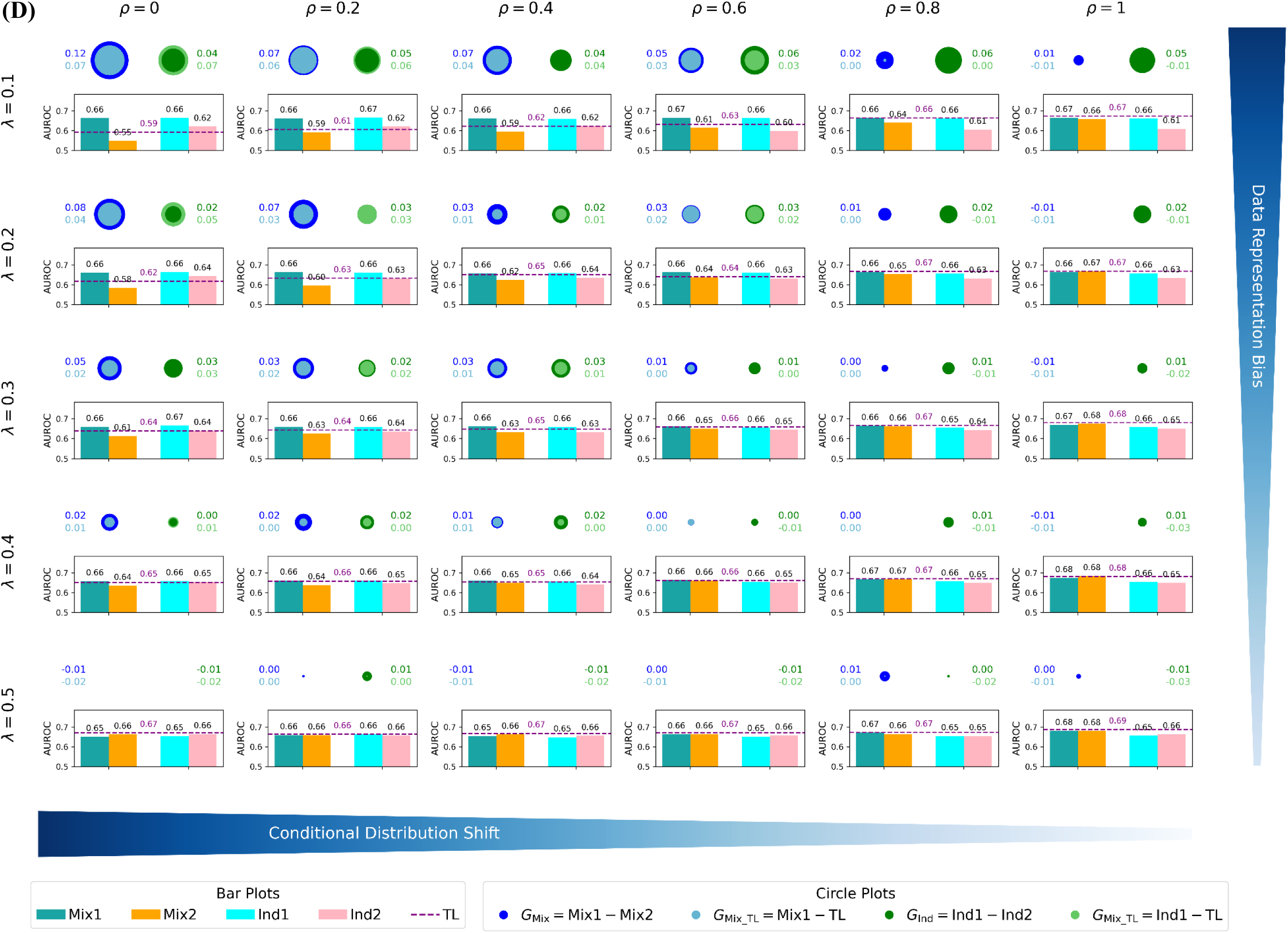
Mapping multi-ancestry machine learning model performance landscapes for the populations consisting of **(A)** EUR and AMR, **(B)** EUR and SAS, **(C)** EUR and EAS, and **(D)** EUR and AFR (*binary* phenotype, h^2^=0.25, number of SNPs=500, total population size=10,000). The hierarchical grid of subplots illustrates the variation in machine learning model performance across a parameter space defined by data representation bias and conditional distribution shift. In each subplot, the bar charts and dashed line show performance of different multi-ancestry machine learning approaches. Mix1 and Mix2 represent the performance of mixture learning for EUR and DDP, Ind1 and Ind2 represent the performance of independent learning for EUR and DDP, and the dashed line represents the performance of transfer learning. The blue and light blue concentric circles represent G_Mix_ and G_Mix_TL_, and the green and light green concentric circles represent G_Ind_ and G_Ind_TL_.

## Notes

### Competing Interest Statement

The authors have declared no competing interest.

### Summary of Updates

Abstract updated to clarify broader relevance to multi-population machine learning beyond polygenic prediction; Introduction expanded to include cross-ancestry prediction methods, nonlinear machine learning, and transfer learning background; Methods section reorganized to precede Results and expanded for reproducibility; Section on multi-population machine learning schemes added to clarify mixture learning, independent learning, and transfer learning as controlled experimental conditions; Synthetic data section updated to clarify genotype coding, effect-size simulation, and rationale for treating SNPs as directly causal; Results section revised to clarify interpretation of λ-ρ performance landscapes and robustness across ancestry pairs, heritability levels, and phenotype types; Discussion substantially revised to emphasize general implications beyond genomics, including dermatology imaging, chest radiograph diagnosis, and education examples; References expanded and reformatted.

